# Doppler-Based Algorithm for Mapping Cardiac Rotors by Induced Temperature Perturbations

**DOI:** 10.1101/362897

**Authors:** Guy Malki, Ofer Barnea, Tamir Tuller

**Author notes:** G. Malki, O. Barnea and T. Tuller are with the Biomedical Engineering Department, Tel-Aviv University, Israel (corresponding).

## Abstract

Electrogram-guided ablation for mapping of abnormal atrial activity has become increasingly popular in clinical applications. However, current methods have several limitations, and none have been shown to increase the ablation procedure success rate more than empirical ablation procedures. Here we present a new approach to identify arrhythmogenic sources as targets for ablation. Based on our previous findings that rotor drifting can be characterized by a local temperature gradient within the tissue, this article describes an innovative induced temperature technique which exploits the fact that rotor drifting produces Doppler shifts in the dominant frequency as measured at stationary locations. A mathematical algorithm is detailed to solve the inverse problem, reconstruct the drift trajectory, and predict the rotor origin location. Mathematical modeling and computer simulations demonstrate the feasibility of the new approach for rotors and focal source, two well-known arrhythmogenic sources of irregular conduction. Performance was extensively investigated for different numbers of electrodes and varied SNRs. Random conditions were also taken into account, since the electrodes’ array position and the initial location of the rotor pivot can impact the outcomes. By using temperature perturbation and employing the Doppler algorithm, the rotor drift trajectory and the origin region is shown to be estimated. We consider ways in which this technique can be extended to differentiate between rotors and ectopic activity. Future experimental and clinical validations should lead to potential use in ablation procedures and improve localization capabilities, thus increasing success rates and optimizing arrhythmia management.

## I. Introduction

A trial fibrillation (AF) is the most common serious abnormal heart rhythm worldwide. It affects more than 10% of the elderly and is the major cardiac cause of stroke [1]. AF is characterized by rapid and irregular activations of the atrium which are often triggered by fibrillatory conduction. This type of conduction is considered to result from one or several stable, self-sustained organized “mother rotors” that can exist in the atria which cause high frequency activation and give rise to the complex patterns of activation that characterize AF [2-3]. Alternatively, AF may be initiated by the existence of focal ectopic sources, mainly in the pulmonary vein - left atrium junction [4].

Rate or rhythm control treatment with antiarrhythmic drugs is considered first-line therapy in patients with symptomatic AF. Yet, pharmacological treatment is frequently ineffective and can have serious potential adverse effects in the long term [5].

For these reasons, surgical ablation procedures have become the preferred treatment for symptomatic AF patients in the last few years. During these semi-invasive procedures, a catheter is inserted into the atria, which delivers radiofrequency (RF) energy that causes local thermal destruction of the suspected arrhythmogenic source areas, and eventually isolates them. This underscores the need to map arrhythmogenic sources correctly during the ablation procedure to both maximize the success of the treatment, as well as to prevent damage through inadvertent thermal injury to the surrounding tissues [6], [7].

In addition, the ablation procedure protocol can vary as a function of the arrhythmogenic source; e.g., ectopic foci vs. rotor, making it crucial to differentiate between them [8].

Several electrogram-guided ablation techniques have been suggested and implemented to address the mapping problem. Most are based on multiple, simultaneously-recorded electrograms that provide spatiotemporal information on the electrical activity in the tissue. These methods are based on analyses in either the time domain, as is the case for the complex fractionated atrial electrogram (CFAE) [9], local activation time (LAT) maps [7], [10], or in the frequency domain such as dominant frequency (DF) analysis [11].

Recently a logical integration of rate and regularity quantities has been developed to achieve computational mapping of the atrial sources [6]. Phase analysis employing phase singularity evaluation has been used to detect rotors and their pivot points [12]. Other complex methods have been suggested to detect the origin points of rotors, including principal component analysis, multi-scale frequency (MSF), kurtosis and spatial Shannon entropy measurement [13], [14]. However, the success rate of these guided-electrogram ablation procedures has not been shown to be better than the empirical ablation procedure, which range from 60 to 90% for both [15]–[17]. In addition, current ablation techniques have other drawbacks such as the ambiguities of the correlations between the underlying arrhythmogenic activity and its electrogram manifestation, and the recurrence of AF after a certain length of time [14], [17]. Thus, finding new methods to improve the detection and the characterization of arrhythmogenic sources during the guided ablation procedures is needed to increase long-term success rates and reduce the unnecessary obliteration of healthy tissue.

It is well known that cardiac spiral wave drifting and meandering can arise without external intervention [18], [19]. Experimental and numerical evidence indicate that rotor drifting towards low excitability regions is caused by spatial ion channel gradients such as the spatial heterogeneity of the I_K_i channel distribution [20], [21] or sodium channel availability [22]. In practice, this drift produces a Doppler-induced difference in local activation periods along the direction of drift [23]. As for any wave source that moves in space, the drifting of spiral waves results in a Doppler shift in the dominant frequency as measured at stationary locations such that the excitation period at a location ahead of the drifting core is shorter than the period behind the core, and the difference is larger for higher drift velocities.

Recently, we showed in a numerical study that spatial heterogeneity in tissue excitability can be induced by local changes in tissue temperature which are achieved artificially by applying a temperature gradient or a regional temperature perturbation (RTP) [24]. We reported that spiral waves drifted towards the colder tissue region associated with the global minimum of excitability. A local perturbation with a temperature of T=28°C was found to be optimal for spiral wave attraction. Unlike the heterogeneity of ion gradients which is a natural phenomenon, a man-made alteration of temperature in a given region of interest can produce a forced controlled drift of the rotor, and create Doppler shifts in different sites within the tissue.

Here, we used 2D numerical simulations to examine clinical applications of this Doppler effect induced by spiral wave drifting after an RTP to estimate the rotor drift trajectory and assess its origin location. This technique can be used to differentiate between spiral waves and ectopic foci since in contrast to spiral waves, ectopic foci are stationary, are not affected by a peripheral temperature perturbation, and do not result in Doppler shifts. This differentiation is of particular importance in both basic science and clinical settings since it will enable experimentalists to determine whether the driving mechanism of atrial fibrillation is a mother rotor or not. Practically it can be employed to optimize ablation techniques since ablation procedures are preferentially carried out for stationary sources (e.g., ectopic foci) whereas drug treatment can be employed to annihilate physiological rotors. This study can thus be harnessed as a proof-of-concept for the design of a new methodology for AF mapping and termination.

## II. METHODS

### A. Biophysical Modeling

Atrial electrical activity in a 30 mm X 30 mm 2D tissue was simulated by numerically solving the following reaction-diffusion partial differential equation under the mono-domain formalism and using the isotropy approximation:

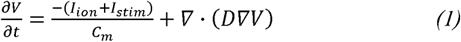

where *V*[*mV*] is the transmembrane voltage, *C*_*m*_[μ*F*/*cm*^2^] is the membrane capacitance per unit area, *I*_*stim*_ and *I*_*ion*_ [μ*F*/*cm*^2^] are the external stimulation and membrane ionic currents, respectively, and *D* [*mm*^2^/*ms*] is the diffusion coefficient that was set to *D* = 0.03 *mm*^2^/*ms* so that a mean planar wave conduction velocity (CV) of ~0.4 m/s was achieved. Eq. (1) was discretized using the finite-difference method in space and the Euler integration method in time, with temporal and spatial resolutions of Δt=2.5 μs and Δh=0.1 mm, respectively.

The chronic AF-modified Courtemanche-Ramirez-Nattel (CRN) kinetic model for human atrial myocytes [25], [26] was chosen for the calculation of *I*_*ion*_, since it is considered to be a robust and simplified-isotropic model that permits activation of a single rotor. Thus, it allows a direct analysis of the relationship between RTP, rotor drifting and the Doppler effect, while avoiding interfering elements such as 3D geometry, tissue anisotropy and heterogeneity, and complex AF conduction patterns that would make the analysis too complex for the proof-of-concept stage.

Temperature effects on the model kinetics were described in detail in our previous publication [24]. Briefly, the rate constants *α*(37°*C*) and *β*(37°*C*) of the gating variables of several ion channels were multiplied by the temperature adjustment factor as in the Arrhenius equation [27] since they have a substantial impact on action potential morphology and dynamics. Specifically, temperature variations lead to significant sustained effects on the action potential duration at 90% repolarization (APD90), as well as the sodium channel recovery time from inactivation, causing both parameters to decrease with increasing temperatures. This observation supports the assumption that low temperature regions in the tissue exhibit reduced excitability, which may eventually lead to a spiral wave drift toward the colder region. Here we used this method to achieve a planned spiral wave drift by employing a local temperature perturbation that resulted in a gradient in the tissue excitability level. The excitability gradient caused a controlled spiral wave drift to the RTP area, which resulted in Doppler shifts that could be used for backward calculation of the rotor location.

### B. Simulation Configuration

All the spiral wave simulations were generated using the S1-S2 cross-field stimulation protocol in a uniform 2D geometry with a fixed temperature of 37°C. First, an S1 stimulation was applied to the left-most column of the tissue (*I*_stim_ = 120*μA*, duration=4ms), followed by an S2 stimulation (*I*_*stim*_ = 40*μA*, duration=3ms) that was applied 160ms after S1 to the top-left or bottom-left quarter of the tissue to initiate a counter-clockwise or clockwise rotating spiral wave, respectively. Spiral waves were generated in the middle of the tissue with a dominant activation frequency of approximately 8.5 Hz. Spiral wave activity was simulated for 5 seconds to guarantee initial stability of the rotor, and only after that stabilization period was a circular 1 mm radius RTP applied. This type of configuration best mimics clinical applications of an external heat source catheter during a guided-ablation procedure [16]. The RTP temperature was set to T=28°C, the optimal temperature to balance out the rapid drift with the stable anchoring around the RTP [24]. The heat distribution due to the usage of a circular heat perturbation was taken into account so that the bio-heat transfer model could be solved numerically [28] in the steady-state for each perturbation configuration using COMSOL finite element software (COMSOL, Burlington, MA).

Our previous simulations revealed that an RTP of T=28°C resulted in an attraction to the colder perturbation characterized by a three-phase pattern: 1) a slow transient phase in which rotors first drift slowly, 2) a fast-transient phase in which the rotors drift rapidly towards the perturbation, and 3) a steady state phase where the rotors anchor around the perturbation. Since the spiral wave moves toward the RTP with a given drifting velocity during the fast-transient phase, Doppler shifts are created at local activations points.

### C. Mathematical formulation of the Doppler-based drifting velocity estimation

In terms of the Doppler effect, the source was defined as the rotor that produces activations at frequency *f*_0_, and the receiver was an electrode placed on the tissue. The activation wave propagates through the tissue at its CV, which is equivalent to the speed of sound in systems that typically use the Doppler effect for sound waves [29]. In our case the receivers were immobile, whereas the source moved at drifting velocity 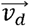. Therefore, the standard Doppler equation was expressed as:

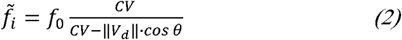

where *f*̃_*i*_ is the obtained frequency at the electrode, ǁ*V*_*d*_ǁ is the magnitude of 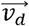, and *θ* is the angle between and the electrode. Eq. (2), however, was adapted for use in the rotor drift scenario. Assume N pseudo-electrodes that are used as observers, which are scattered in a circle with equal internodes of *dθ* = 2*π*/*N* [Fig. 1(a)]. Each local electrode records the electrical signal at its location, as well as the local tissue temperature. *θ*_*d*_ defined as the drift direction; e.g., the angle between 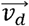 and the vertical direction [Fig. 1(b)]. Therefore, the angle between the i-th electrode and 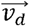 can be written as:

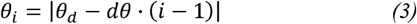

Thus, for N electrodes, we can write N Doppler equations:

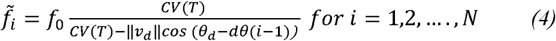

This non-linear equation has three unknowns (*v*_*d*_, *θ*_*d*_, *f*_0_), and can be solved by using such non-linear numerical methods as the Newton-Raphson method. A simpler way would be to solve the equations using linear algebraic methods that eliminate the cosine function by using the definition of the dot product. Since *θ*_*i*_ is the angle between two vectors (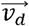 and 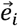, which is the location of the i^th^ electrode) we can write:

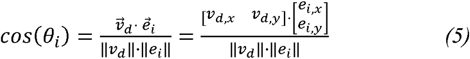

Thus [*v*_*d*, *x*_, *v*_*d*, *y*_] and [*e*_*i*, *x*_, *e*_*i*, *y*_] are the Cartesian components of 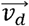 and 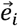, respectively, and ǁ*v*_*d*_ǁ, ǁ*e*_*i*_ǁ are their absolute values. Consequently, (4) can be written as:

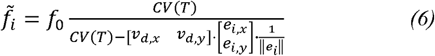

This equation can be re-arranged as a linear equation for (*v*_*d*, *x*_, *v*_*d*, *y*_, *f*_0_):

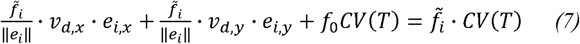

or in matrix form:

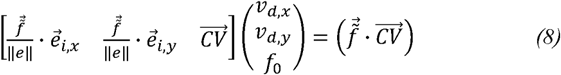

where *f*̃_*i*_, ǁe_*i*_ǁ, *e*_*i*, *x*_, *e*_*i*, *y*_, and *CV*(*T*) for each of the electrodes are stored in vector form as 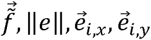 and 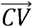.

**Fig. 1.**
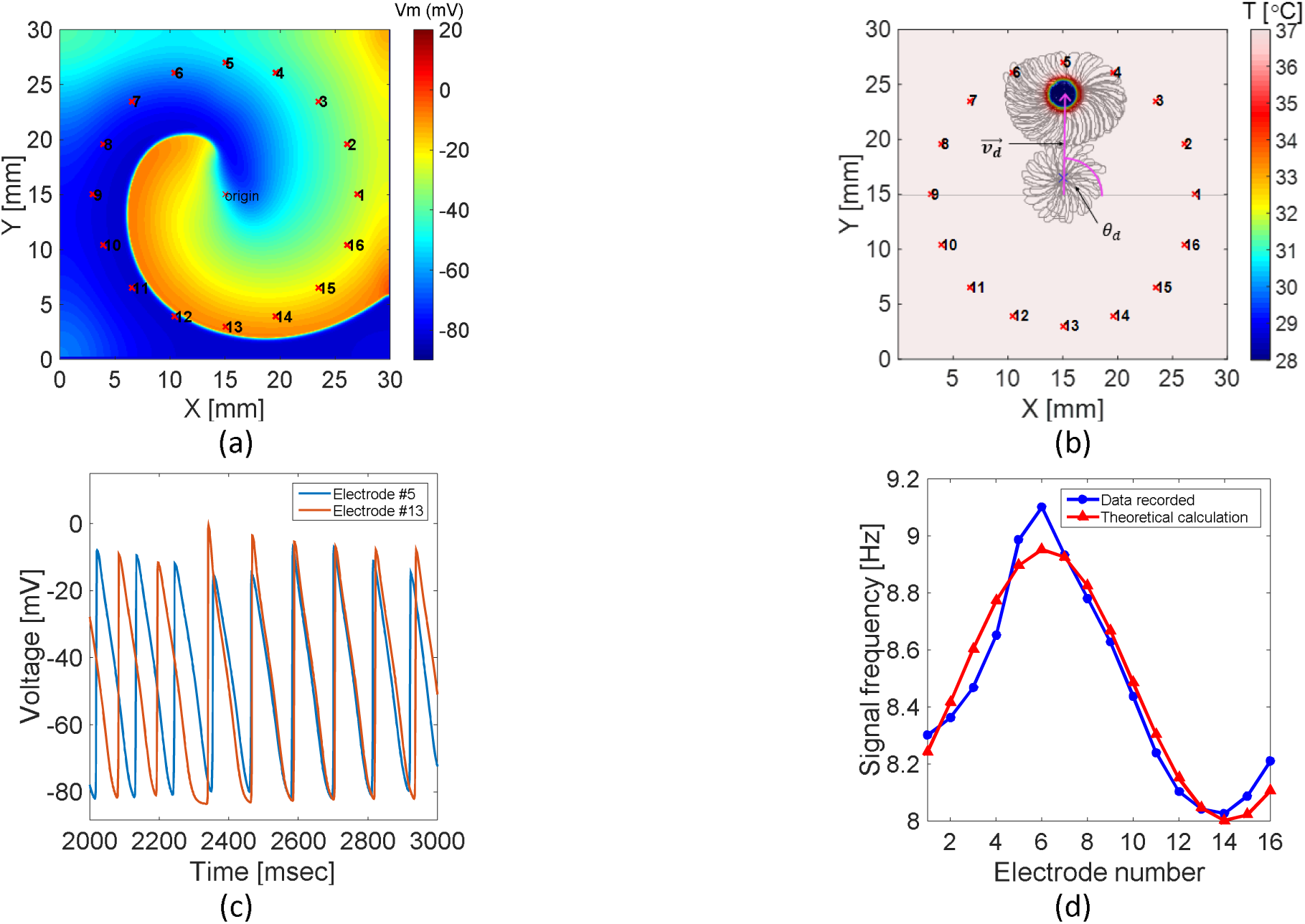
Inputs to the Doppler based reconstruction algorithm, (a) snapshot of a spiral wave in a 2D tissue, indicating its origin location in the middle, (b). Temperature map with RTP of T=28°C. The spiral wave tip trajectory is plotted as a gray line, and demonstrates the drift from the origin location in the center toward the RTP, ending in anchorage around it. The drift velocity vector (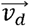) and the drift angle (*θ*_*d*_) are shown in the figure. Both A and B display a 16-pseudo electrode array, placed in a symmetric circular configuration with a radius of 12.5 mm. (c). Two recorded signals from electrode #5 and electrode #13 at the window time of 2000-3000 ms, which is the fast-transient phase in which the rotors were rapidly drifting towards the perturbation in response to the RTP in B. The difference in the signal frequencies according to the relative location of the electrodes from the drifting direction, (d). Frequency analysis using FFT for all the electrode arrays for the above time segment depicts the spectral differences between the signals (blue). Theoretical calculation of these frequencies using the Doppler equations for a known drift velocity (red) reveals the small bias that arises from the spiral motion which reduces by using a window analysis.

This set of the Doppler equations was validated by simulating a moving point source at T=37°C with an activation frequency of 5Hz, and a known velocity of the form 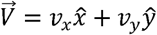. The simulations were run for several combinations of v_x and v_y, so that 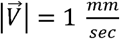, and the direction of motion relative to the horizontal axis was varied from zero to 2π, at 22.5° increments, yielding 16 configurations. All simulations lasted 10s with a sampling frequency of 1 kHz, and included 16 pseudo-electrodes for signal recordings that were scattered in a circular shape with a radius of 12.5mm. Drift velocity and activation frequency were computed according to (8), and were compared to their real values at each simulation.

### D. Reconstruction algorithm

Using the above method, an algorithm to reconstruct the spiral wave drift trajectory and decipher its origin point was developed. An array of electrodes was created by determining their spatial arrangement, positions and labels. In all simulations 16 pseudo-electrodes were used, since this is a power of two that represents a symmetric distribution on the tissue in which the spatial symmetric records contribute to a better understanding and analysis of the results at this stage. For each of the electrodes a 1D voltage signal as a function of time with a sampling frequency of 1 kHz was recorded. Fig. 1 depicts the representative voltage (a) and temperature (b) maps with the array of 16 electrodes that was described previously. An illustration of a spiral wave drift trajectory towards and anchored around the cold perturbations appears in Fig. 1(a).

To obtain the dominant frequency vector, 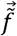, an FFT method with a frequency resolution of 0.01 Hz was implemented for the multiplication of each signal with a symmetric Hamming window. The example in Fig. 1 panel C shows signals from two electrodes during the rapid drifting phase. Electrode 5 is located beyond the RTP; hence the rotor is drifting towards it, whereas electrode 13 is situated in the opposite direction and the rotor is receding from it. Clearly, the signal in electrode 5 has more activations per time window than electrode 13; thus, its frequency is much larger (8.98 Hz. Vs. 8.04 Hz for electors 5 and 13, respectively). The blue line plotted in Panel D depicts the frequencies of all the electrodes at the above time window. It shows that signals that were recorded in electrodes located where the spiral wave drift was approaching them had higher frequencies than the electrode signals where the spiral wave was moving away from them.

The values of *e*_*i*, *x*_, *e*_*i*, *y*_ were assigned to each electrode based on its coordinates, and the electrode’s norm was calculated as 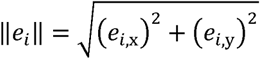. The matrix columns that appear in (8) 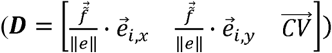 defined and the solution vector 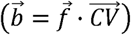 was formed. Using the QR-factorization method the linear system 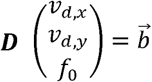 was solved.

In contrast to a moving point-source that progresses in one direction at all times, a spiral wave has a more complex movement that influences the Doppler effect. When espousing the rotational movement, the rotor core changes direction constantly, but unlike a point source, it does not advance to the colder region at each time point. The net drift is toward the colder area, but there are phases where the core propagates to other directions due to its spiral nature. Consequently, there is a small bias between the frequencies obtained for the spiral wave when using this method and the calculated theoretical Doppler frequencies for a known drift velocity and activation frequency, as depicted by the red line in Fig. 1 Panel D. This problem can be significantly reduced by using a time window analysis. The ideal window in this case is equal to the tip rotation cycle (e.g. analysis every 1000 ms); hence the data were sampled in roughly similar phases, with an overlap of 50% between two successive windows. This is considered to be an optimal overlap value that corrects for the bias from the spectral estimation and its variance. Therefore, for N electrodes and M time segments, the analysis covered N by M signals.

The next stage involved the spiral wave trajectory reconstruction using the estimated velocity components at each time window. As stated above, a local small temperature perturbation can be employed to attract and anchor a distant spiral wave after a short time. Since the spiral wave is anchored around the perturbation we created, its final location (denoted as [*x*_1_, *y*_1_]) is known. By using the time window velocity vectors, a reconstruction of the spiral wave displacement vectors 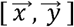 from the endpoint to its origin point can be computed at each Δt time frame as follows:

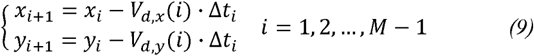

As presented in the Results, the reconstructed trajectory required a shifting and rotating correction to improve its accuracy. The first step involved the detection of the time steps making up the anchoring phase. This was done by identifying the time segment that had the largest change in displacement either in the X or Y components. This time segment was associated with the fast-transient phase in which the rotors drift rapidly towards the perturbation, and was defined as the critical time step. Thus, all the previous time steps in the reconstructed trajectory were associated with the steady state phase where the rotor was anchored around the perturbation. Next, the center of mass of the estimated trajectory in the steady state time step phase was computed, which yielded the estimated perturbation center. This was compared to the known location of the perturbation. The difference was defined as the fixed vector 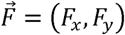. By knowing 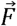 we could shift the whole trajectory so the anchoring pattern would be around the known perturbation. In addition, we calculated the angle of rotation relative to the perturbation location *φ* = tan^–1^(*F*_*y*_/*F*_*x*_), and hence could rotate the trajectory by using the 2D rotation matrix:

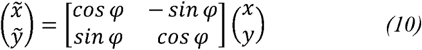

### E. Algorithm performance evaluation

The real tip trajectory of the drifted rotor, which served as the gold standard, was automatically tracked as in Nayak *et al*. [30]. The mean trajectory at each tip rotation cycle was calculated to obtain 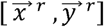. Then, to quantify the algorithm error, the mean standard error (MSE) between the real and the reconstructed trajectories was calculated for each component as:

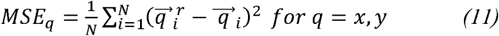

This yields the total MSE:

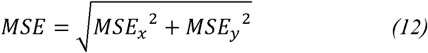

To investigate the algorithm characteristics and to optimize its performance, several parameters were explored. First, the reconstruction algorithm was run with 3 to 64 electrodes, and the total MSE was measured as a function of the number of electrodes. Then the performance of the algorithm was examined in noisy environment by adding white Gaussian noise to the signals, with a signal-to-noise ratio (SNR) from 5 dB to 40 dB, in 5dB increments. Twenty simulations at specific SNR values and specific numbers of electrodes were conducted (for a total of 20 X 9 X 62= 11160 simulations), and the mean total MSE value for each set was calculated.

Asymmetric arrangements of the electrode array relative to the 30 mm X 30 mm tissue size were examined by changing the center of the circular electrode array. Thus, the X and the Y coordinates of the circular array’s center were modified from 5 to 25mm. As a function of this shift some of the electrode locations were located outside the 2D tissue and were removed, such that only the electrodes that were within the 30X30 tissue were used in the algorithm.

### F. Differentiating between the rotor and focal source

One goal of this study was motivated by the need to differentiate between focal to reentry sources. Two types of arrhythmogenic drivers were simulated: rotors by using standard S1-S2 cross field stimulation, and focal activity using S1 point stimulations with a similar frequency as the rotors (typically 8.5Hz in the atrial tissue). For each type of source, 200 different simulations were performed, which differed from each other in both the arrhythmogenic source location and the position of the electrode array. Both parameters were randomized in each simulation. This protocol was designed to simulate cases that can occur in real clinical setup. Activation frequencies were measured at each simulation, and the spectral differences were analyzed. Using these results, a decision process was implemented to determine whether the arrhythmogenic source was ectopic or reentry. Afterwards, for all the spiral wave simulations, the reconstruction algorithm was applied to analyze the algorithm results for arbitrary conditions, and the distance between the real and the predicted origin point was computed and compared.

### G. Technical specifications

All simulations were performed using C++ code running on a high-performance cluster computer (Altix X86-PTO; Silicon Graphics International, Milpitas, CA) with a master node (8 cores, Xeon 2.5 GHz processor; Intel, Santa Clara, CA) and up to five computational nodes (60cores, Xeon 2.8 GHz; Intel). Data analysis, the reconstruction algorithm and visualization were performed with MATLAB R2014b (The MathWorks, Natick, MA).

## III. Results

### A. Model validation

The Doppler equation system was validated with a moving source with 16 different configurations of velocity vectors. An activation frequency of 5 Hz on a 30 mm X 30 mm atrial tissue model using the CRN model with chronic AF modifications was used. Fig. 2(a) shows the 16 real and estimated velocity vectors, depicted as blue and red lines, respectively. Each configuration is numbered in terms of its related vectors. For example, configuration 1 represents a velocity vector with 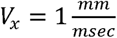 and 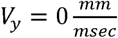, whereas the configuration 2 components are 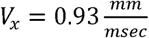 and 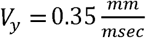. As can be seen, the algorithm evaluated the true velocity values correctly, with a total MSE of 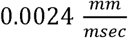.

**Fig. 2.**
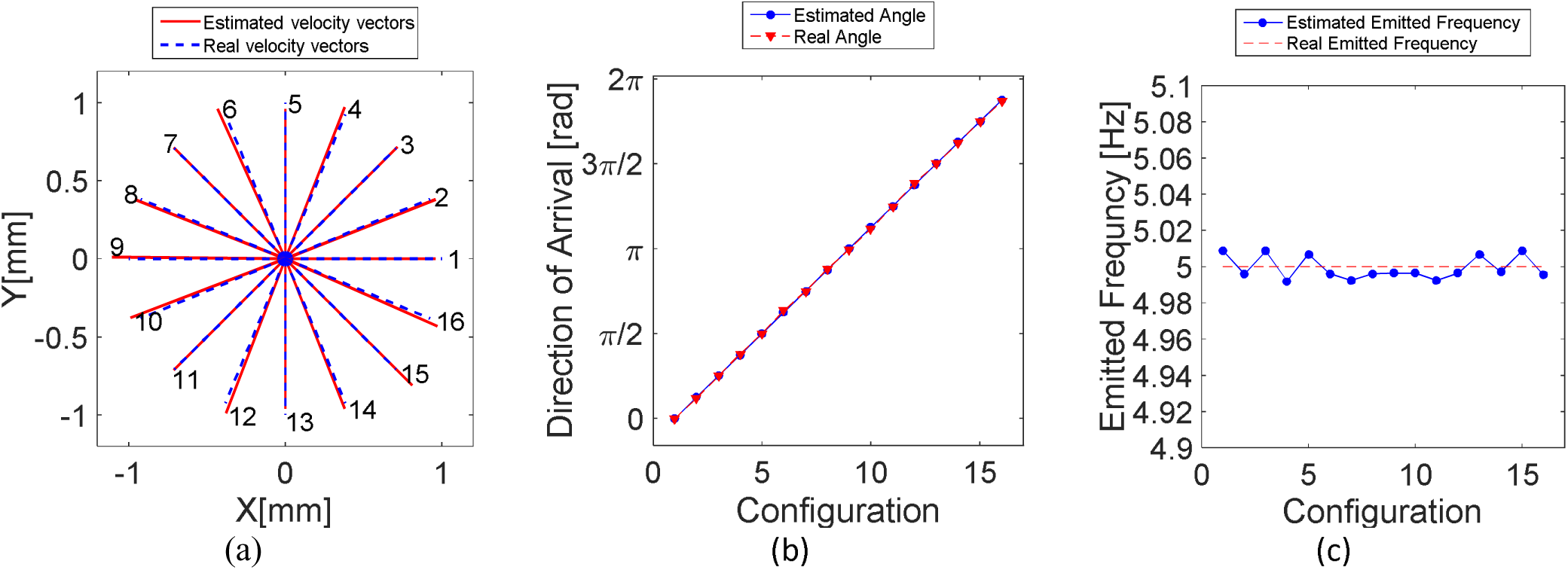
Validation of the Doppler equation system: Simulations of a moving point source with an emitting frequency of 5 *Hz* was conducted with 16 combinations of *V*_*x*_*x*̂ + *V*_*y*_*y*̂ = 1 at increments of 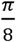 between each consecutive vector. The Doppler equation system was applied to estimate the velocity vector (a), the direction of arrival (b), and the activation frequency (c) of the moving source. In all panels the estimated values (blue) are plotted against the real values (red), for each configuration.

Vectors that are parallel to the X or Y axis (1, 5, 9 and 13), or at an angle of 45° ± 90°*n* relative to these axes (3, 7, 11 and 15), had much better results with a total MSE of 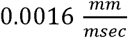, whereas in the other directions the total MSE was 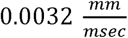. Since the configurations were dependent on the angle between *V*_*x*_ and *V*_*x*_, we were also able to investigate the direction of movement, as shown in Fig.2(b) where the estimated angle is similar to the real angle, with a mean error of 0.68° ± 0.47° and a maximum error of 1.7°. The third variable of the linear system, the activation frequency, was also calculated and compared to the original rate of 5 *Hz*, as displayed in Fig.2(c). The mean error was 0.0058 *Hz* + 0.002 *Hz*, with a maximal error of 0.0086 *Hz*, which is considered extremely small. These observations indicate that the mathematical formalism of the Doppler equations system is reliable for these 16 possible cases of directions, which match different combinations of negative or positive for both the X and Y components, and varying ratios between *V*_*x*_ and *V*_*y*_. Thus, this method can be used for the complete trajectory reconstruction algorithm. However, the dissimilarities between a focal source and a spiral wave should be taken into account, as discussed below.

### B. Reconstruction of spiral wave trajectory algorithm

The drift velocity vector at each time frame was estimated beforehand using the Doppler linear equation system (8), and the drift displacement vectors at each time segment were calculated (9). A typical example is shown in Fig. 3, using the RTP displayed in Fig. 1(b) which is centered at (*x*_1_ = 15 *mm*, *y*_1_ = 24 *mm*), and an array of 16 electrodes as illustrated in the figure. Panel A displays the reconstructed drifted line for the whole tissue (left), and a zoomed-in view of the part of the tissue where the drift trajectory occurred (right). Each displacement vector is colored differently to represent a specific time frame. It is plotted as an arrow that points to its backward direction, corresponding to the backward calculation, ending with the estimation of the rotor origin, which stands for the front end of the last arrow. The estimated backward trajectory is shown to adhere to an initial circular route equivalent to the steady state anchoring phase around the perturbation, and is followed by a quick propagation for two time frames toward the estimated origin point, which is considered as the fast-transient phase. The black dashed line, separated by triangles, is the real calculated trajectory divided into the same time segments. Panel B compares the real (red) and the estimated (blue) backward trajectories for the X (left) and the Y (right) components.

**Fig. 3.**
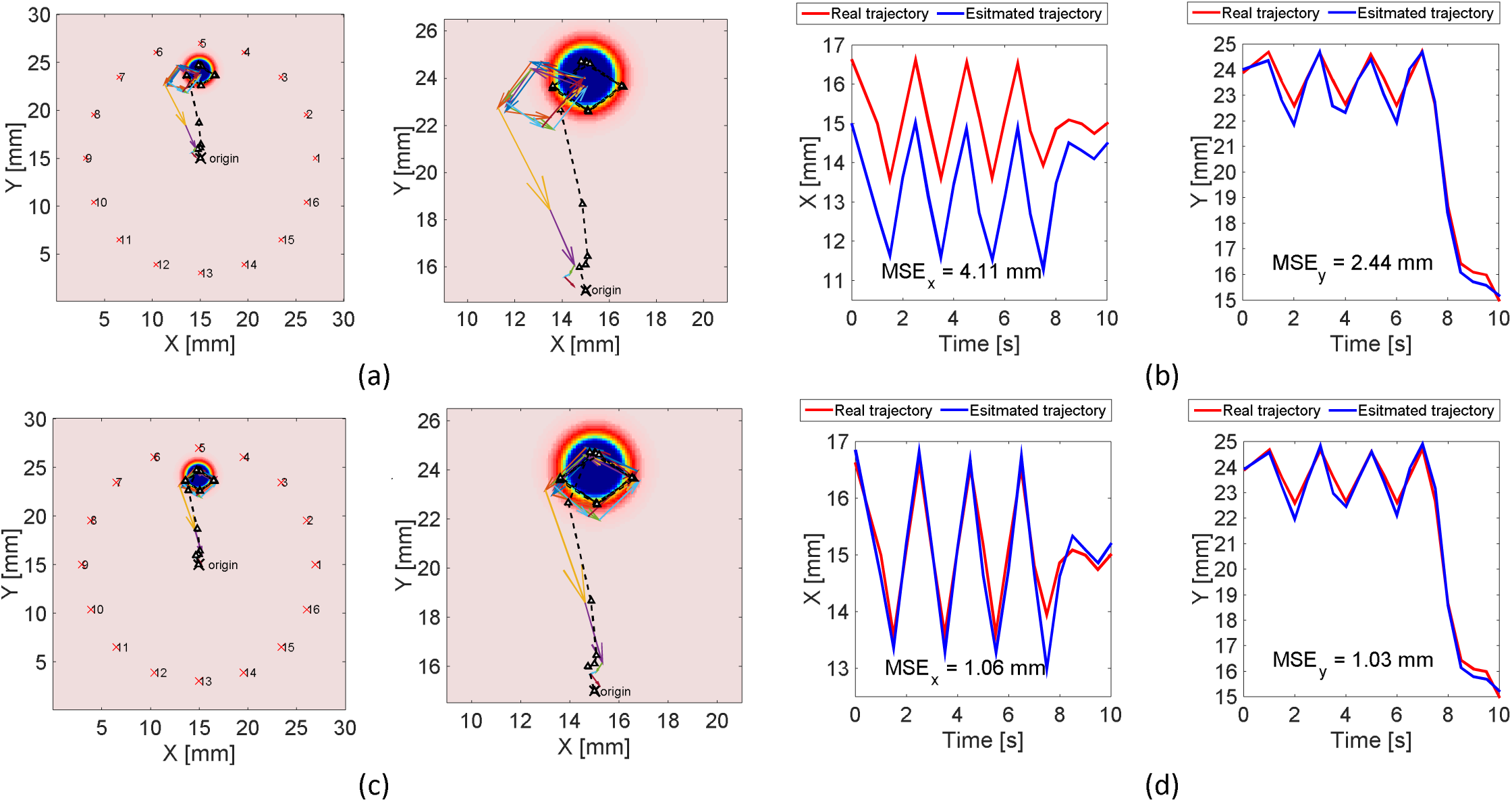
Reconstruction of the drifted spiral wave trajectory before (top) and after (bottom) correction. The estimated drifted line is plotted onto the whole tissue (left) and with a zoomed-in section (right), where each arrow represents the reconstructed displacement vector for a specific time segment and is shown in a different color. The original mean trajectory is plotted as the dotted black line using the same time segments (a, c). Comparison of the real (red) to the estimated (blue) trajectories as a function of time for the X (left) and the Y (right) directions. The MSE for each component is shown (b, d).

There was a successful match concerning the behavior of the reconstructed route: at any given moment the pattern of the estimated trajectory was in the same phase as the original trajectory, and the peak-to-peak amplitude was also similar. However, the total MSE was 4.78 mm, thus presenting a discrepancy between the real and estimated coordinates, which yielded an erroneous trajectory, and ultimately an incorrect evaluation of the rotor origin point. Therefore, this evaluation was corrected by a shift and rotation of the trajectory, as described in the Methods. The reconstructed trajectory is plotted in Fig. 3(c), and the real and estimated coordinates for the X and the Y components are compared in Fig. 3(d), similar to the previous illustrations. The results indicate a significant improvement in the reconstruction, that reduced the MSE_X_, MSE_Y_ and the total MSE by 75%, 58% and 69% to 1.06 mm, 1.03 mm and 1.42 mm, respectively. The distance between the estimated origin point of the rotor to its real one was less than 3 mm in this case.

The performance of the algorithm was also assessed by varying the number of electrodes (N) in the circular array from 3 to 64, with an equal gap between each adjacent pair. The minimum number of electrodes was 3, because the Doppler equations system is solved for 3 variables. Fig. 4(a) presents the total MSE as a function of the number of electrodes. As expected, MSE decreased as the number of electrodes increased, since there is more information that contributes to the optimization of the results using the QR factorization calculation. The MSE showed an exponentially decreasing dependence on N, in terms of the best-fit relationship (R^2^=0.9991),

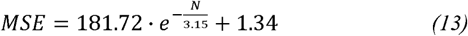

Thus, the least number of electrodes required for a minimal total MSE of the reconstruction algorithm is 20 electrodes, since increasing N beyond this value had no further effect on the MSE. In addition, the algorithm performance was examined in a noisy environment by adding a white Gaussian noise to the signals, with a SNR ratio of 5 dB to 40 dB. As shown in Fig. 4(b), simulations with a SNR of 20dB and up led to the same MSE. However, the simulations confirmed the exponentially decreasing relationship between the number of electrodes and the MSE. To underscore the dependency of the MSE on both the SNR and number of electrodes, graphs of MSE as a function of a SNR for 12, 16, 32, 48 and 64 electrodes are shown in Fig. 4(c). This illustration trivially shows that as the SNR increases, the MSE decreases. More specifically, at each specific SNR, as more electrodes are used the MSE drops. Nevertheless, these changes become increasingly smaller as the SNR rises. Clearly, in all configurations except for the 12 electrode array, the MSE becomes unified for a SNR above 30 dB.

**Fig. 4.**
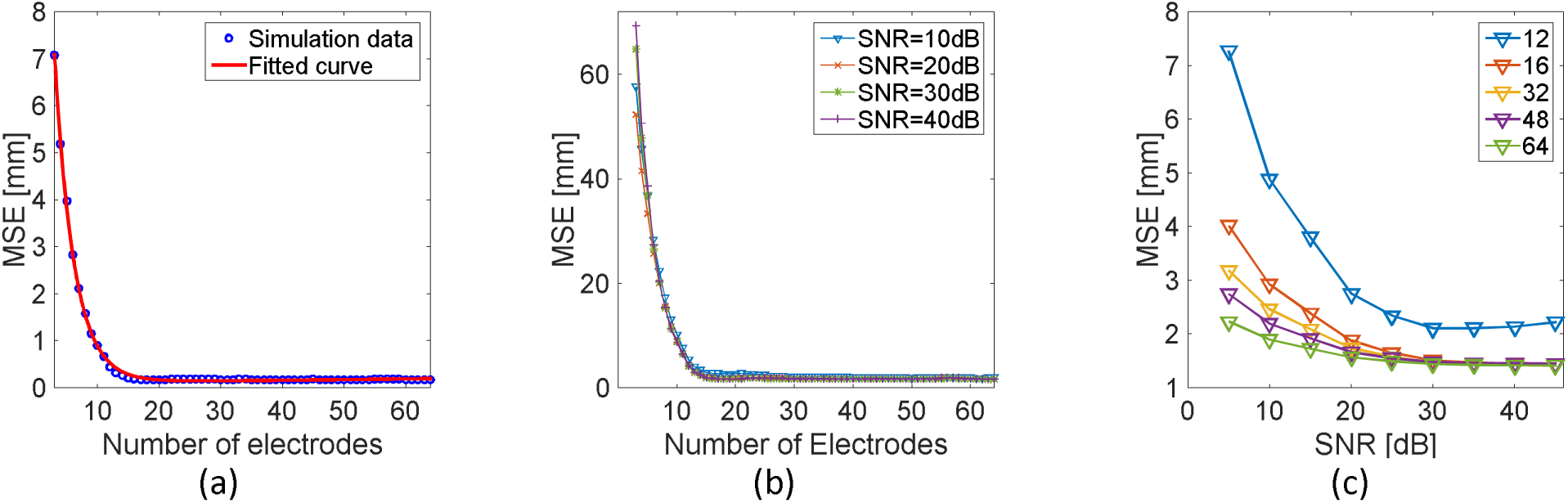
The number of electrodes and SNR influences the algorithm performance. MSE is plotted as function of the number of electrodes (blue circles) and an exponentially decreasing curve was fitted with R2=0.9991 (a). Same analysis for a varying SNR (b). MSE as a function of the SNR for different numbers of electrodes (12, 16, 32, 48, 64) demonstrated exponential behavior (c).

Next, we investigated the algorithm outcomes using asymmetric electrodes arrays; e.g., when the center of the circular array was not located on the rotor origin. For this purpose the X or Y coordinates of the electrode array center were varied from 5 to 25 mm, yielding 441 combinations. Fig. 5 illustrates several representative examples of the estimated trajectories, where X and Y varied separately up to 16, 20 and 24 mm. It should be noted that according to the array shifts, some electrodes were located outside the tissue and therefore were omitted, resulting in fewer signals for the backward calculation. However, as can be seen, in most cases the trajectory was assessed correctly, and the rotor origin area could be identified properly.

**Fig. 5.**
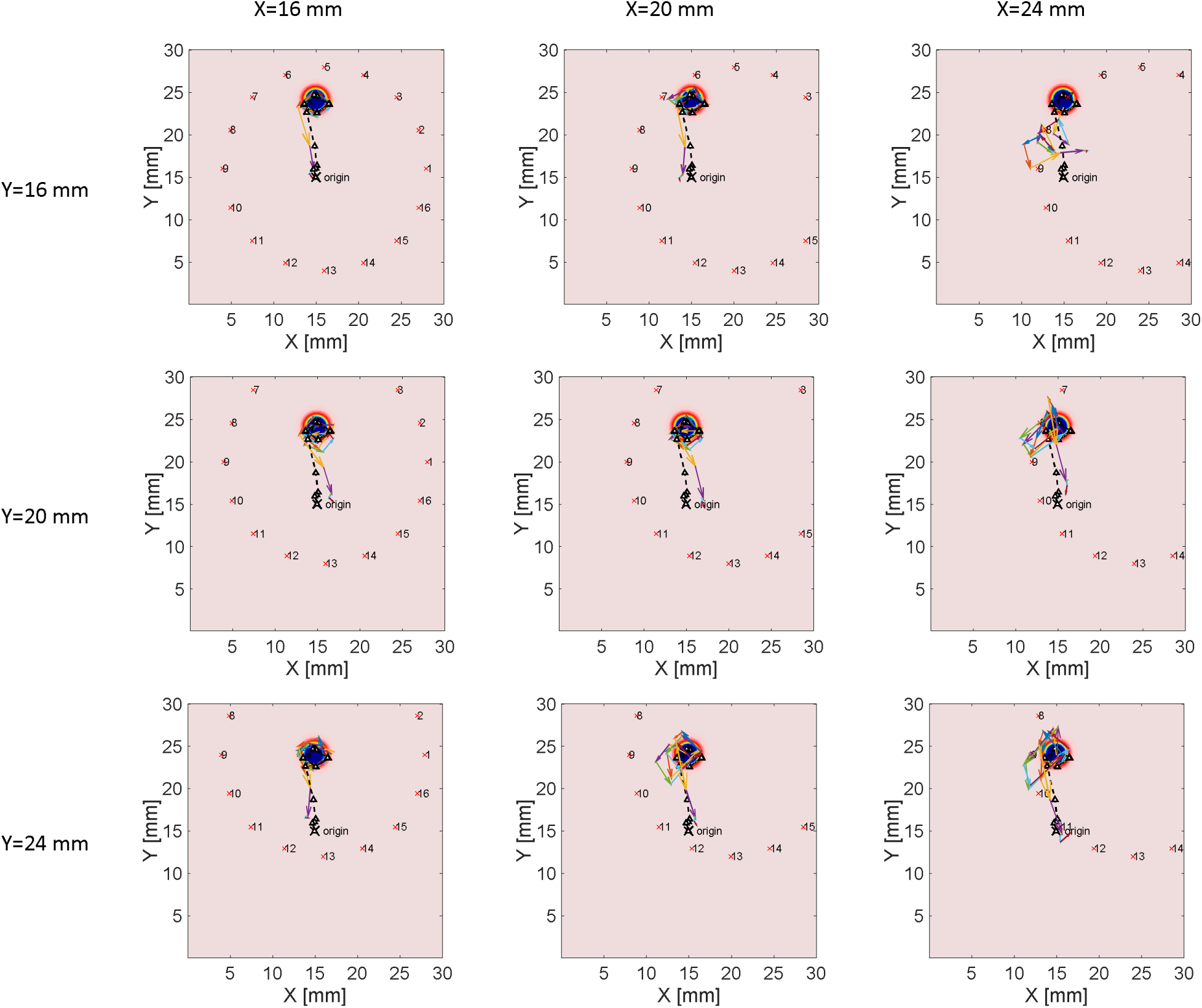
Algorithm results for asymmetric positions of the electrode array. Nine examples are shown in the figure, when each component of the center of the electrode array was set at 16, 20 or 24 mm. The estimated trajectory is plotted with colored arrows, and the real trajectory is plotted with a black dotted line. Each subplot shows the RTP, and the rotor origin point marked by ‘X’.

A more general analysis of all simulated arrangements is depicted in Fig. 6(a). The bottom layer indicates the 2D temperature map including the RTP and the origin location of the rotor (marked by X). On top, a square is drawn in which each pixel represents the location of the center of the electrode array for different simulations, and the MSE for each case is color-coded. Overall, as the circle of electrodes contains both the RTP and the rotor origin point within it the reconstruction is satisfactory. Hence, the MSE is low when the electrodes center is closer to the rotor origin and stays small as the Y coordinate is varied. Nonetheless, displacements of the X coordinate to the outermost locations of the X axis cause the MSE to increase rapidly.

**Fig. 6.**
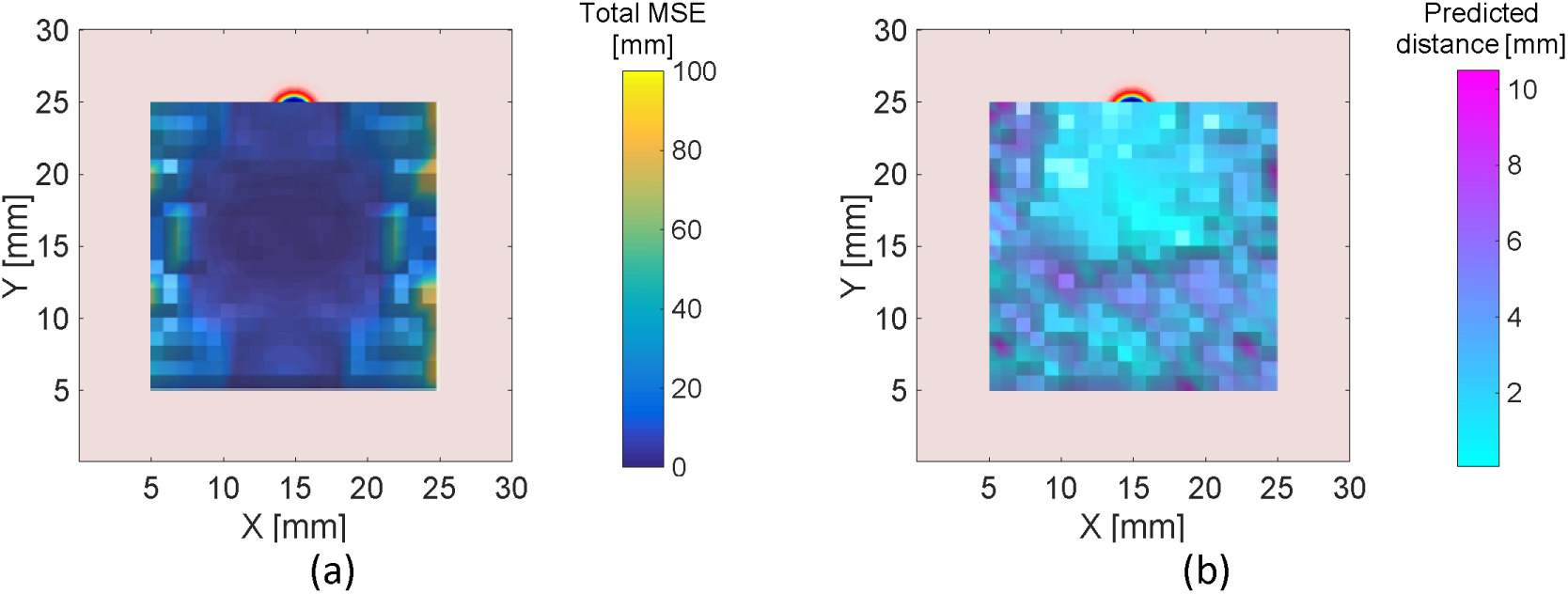
Analysis of the algorithm results for asymmetric positions of the electrode array. Each panel shows the temperature map in the background, and a 2D map where each pixel represents the center of the electrode array, and its color indicates the total MSE (a) or the predicted distance between the estimated and the real origin point (b).

This observation can be explained by the fact that less information reaches the electrodes since some electrodes were excluded. In addition, when an electrode is fortuitously placed exactly on the RTP, as in the case of X=24mm, Y=16mm in Fig. 5, the backward calculation does not work well. In addition, when the electrodes become distant from the drift route, the Doppler effect is much less noticeable, making the algorithm less efficient. Nevertheless, this scenario can be avoided since the electrode array and the RTP locations are controlled by the user and can be designed appropriately. In all other cases, even if the foreseeable drift line is not accurate enough and cause the MSE to increase, the predictable region of the rotor origin is reliable within a small margin of error. To quantify this outcome, a similar plot is illustrated in Fig. 6(b) in which the distance between the estimated and the real rotor origin point is the color-coded variable. In the majority of cases the predicted origin location can be estimated close to the original one; thus, the distance between the algorithm’s predicted end-point and the actual rotor origin location is at most 9.7mm with an average value of 2.68 mm ± 1.4 mm.

### C. Differentiating between rotor and focal sources

Simulations of spiral wave and focal activity in the presence of RTP were carried out for 200 different arbitrary combinations of origin point and electrode array positions for each arrhythmogenic source. Characteristic snapshots of the electrical waves can be seen in Fig. 7(a) for both types, along with the randomly located electrode array. In each simulation 16 electrodes recorded the electrical signals, and spectral analysis was performed to obtain the dominant frequency in each electrode, as detailed above. Fig. 7(b) presents the activation frequencies at each electrode for the focal source and the spiral wave, respectively, which appear in panel A. As can be inferred from these representative plots, the spectral signal of the focal source is constant up to negligible numerical errors. On the other hand, the drifted rotor led to variance in the activation frequencies, as expected. These results are a direct consequence of the Doppler effect in the presence of drifted rotor, which does not occur in the case of focal activity, supporting the assumption that a rotor can be distinguished from focal source by employing a local temperature perturbation.

**Fig. 7.**
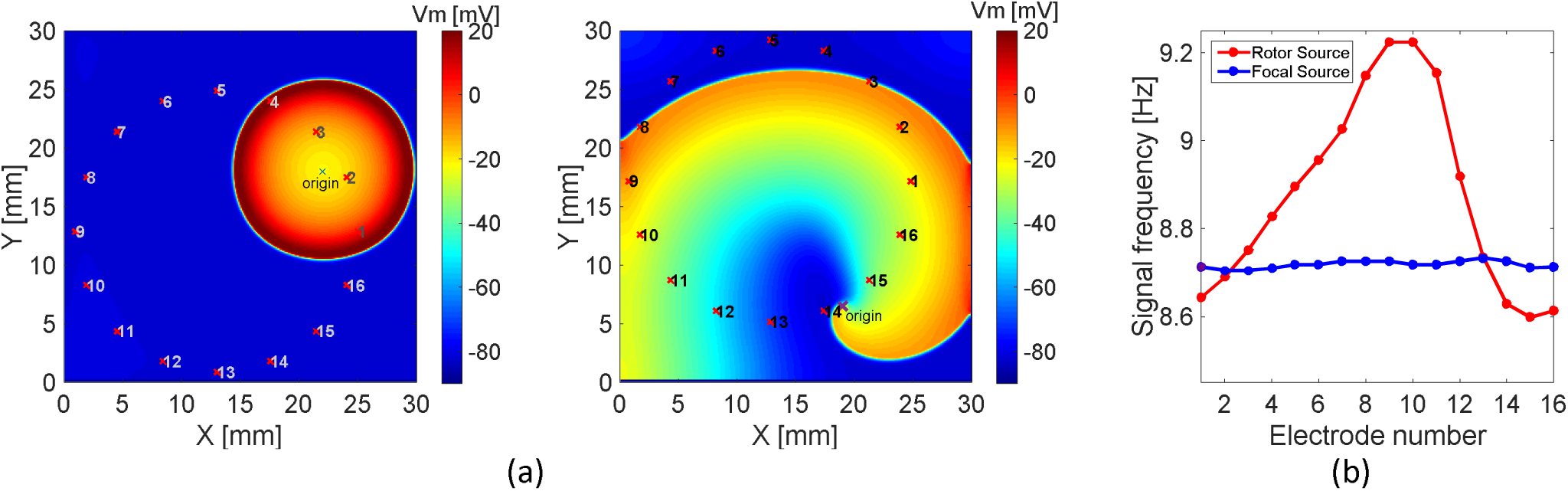
Spectral differences between an ectopic source and a reentry source. Typical voltage map snapshots of a simulated focal source (left) and a spiral wave (right), respectively, which were initiated at an arbitrary origin point in the tissue. In each simulation 16 electrodes were placed in a circular array with a random center (a). The dominant frequencies of the signals of each electrode were calculated and plotted (b). The spiral wave exhibited a major difference in frequencies due to the Doppler effect, as expected. However, the computed frequencies for the focal source were a constant value, yielding a clear difference between the two sources.

The practical distinction between these two source types based on the spectral data [Fig. 7(b)] can be made by applying simple mathematical tools. We computed the variance of each frequency vector, since the variance in the case of focal activity is expected to be zero. In addition, using the variance reduces the impact of outliers that can occur if the RTP or the electrode is somehow located exactly on the point source. Thus, a small threshold can be set to ensure a binary decision as to the presence of a focal source or a spiral wave. If a spiral wave is detected, the reconstruction algorithm can be employed to detect the rotor origin’s location. The complete flowchart for this process is presented in Fig. 8.

**Fig. 8.**
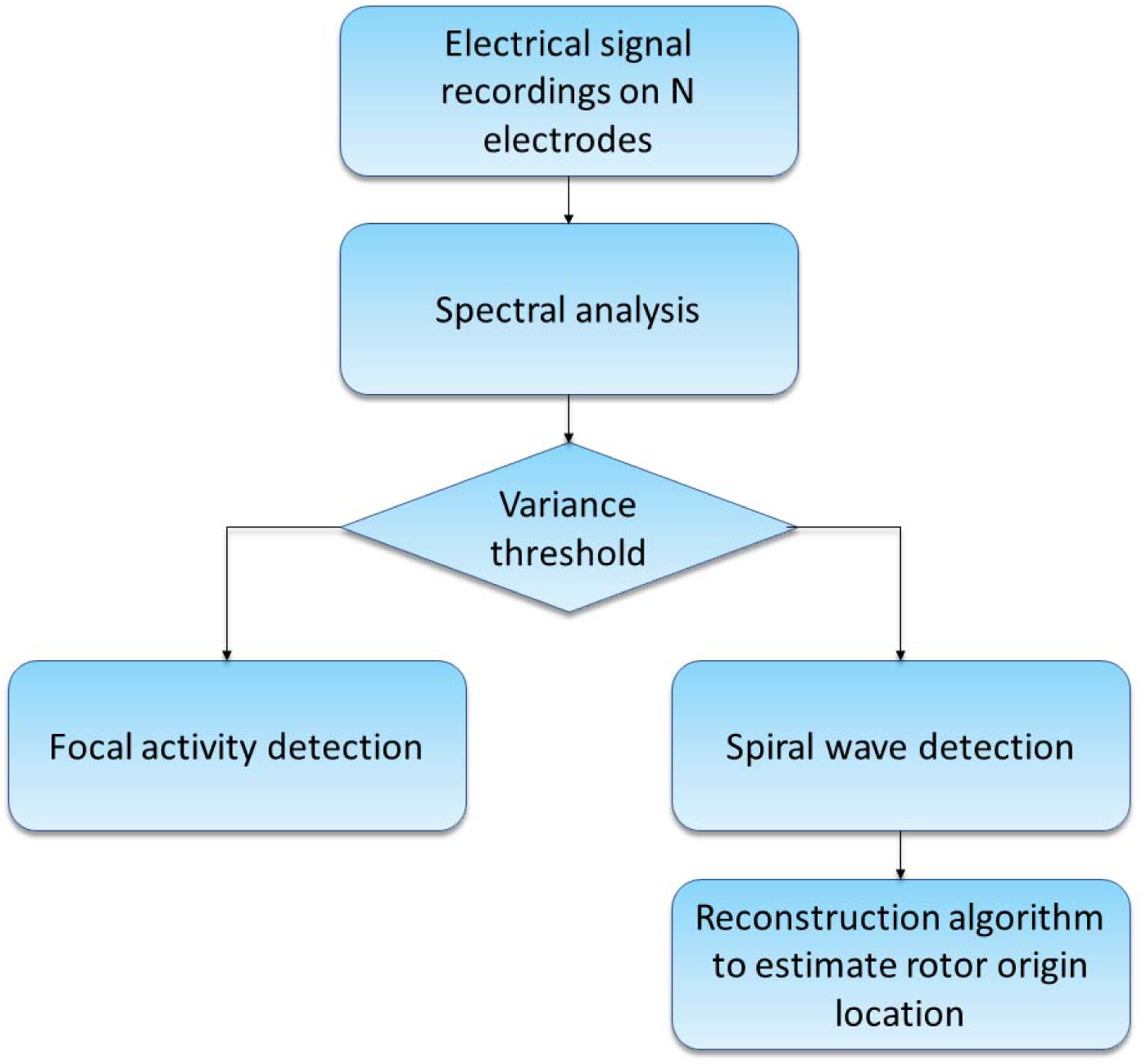
Full algorithm flowchart diagram. The algorithm inputs are multiple electrical signals from the atrial tissue. A frequency analysis is performed on these data and using the spectral variance, a binary decision for the detection of a focal source or rotors is made. In the case of rotor detection, its activity area is estimated using the Doppler-based reconstruction algorithm.

The full process was implemented on the 400 simulations. For each simulation the variance of the frequency vector was calculated. Fig. 9 shows the histograms for the frequency vector variance for each focal activity (blue) and spiral wave (red) simulation. This diagram illustrates the unequivocal distinction between the two samples, and clearly identifies the threshold of the frequency vector variance to be between 0.016 and0.05 Hz. Fig. 10 shows a typical reconstruction of the spiral wave trajectory and detection of the origin point for the rotor shown in Fig. 7(a). This plot is an example of the success of the algorithm to locate the origin area in asymmetric conditions and random sites, as would be expected in real scenarios. Using this method, all the 200 ectopic foci simulations were identified correctly as a point source. The 200 random spiral wave simulations were also detected properly as a rotor source although the origin point detection using the reconstruction algorithm had slightly different results. A histogram of the Euclidean distance between the real and the estimated origin point is presented in Fig. 11(a).

**Fig. 9.**
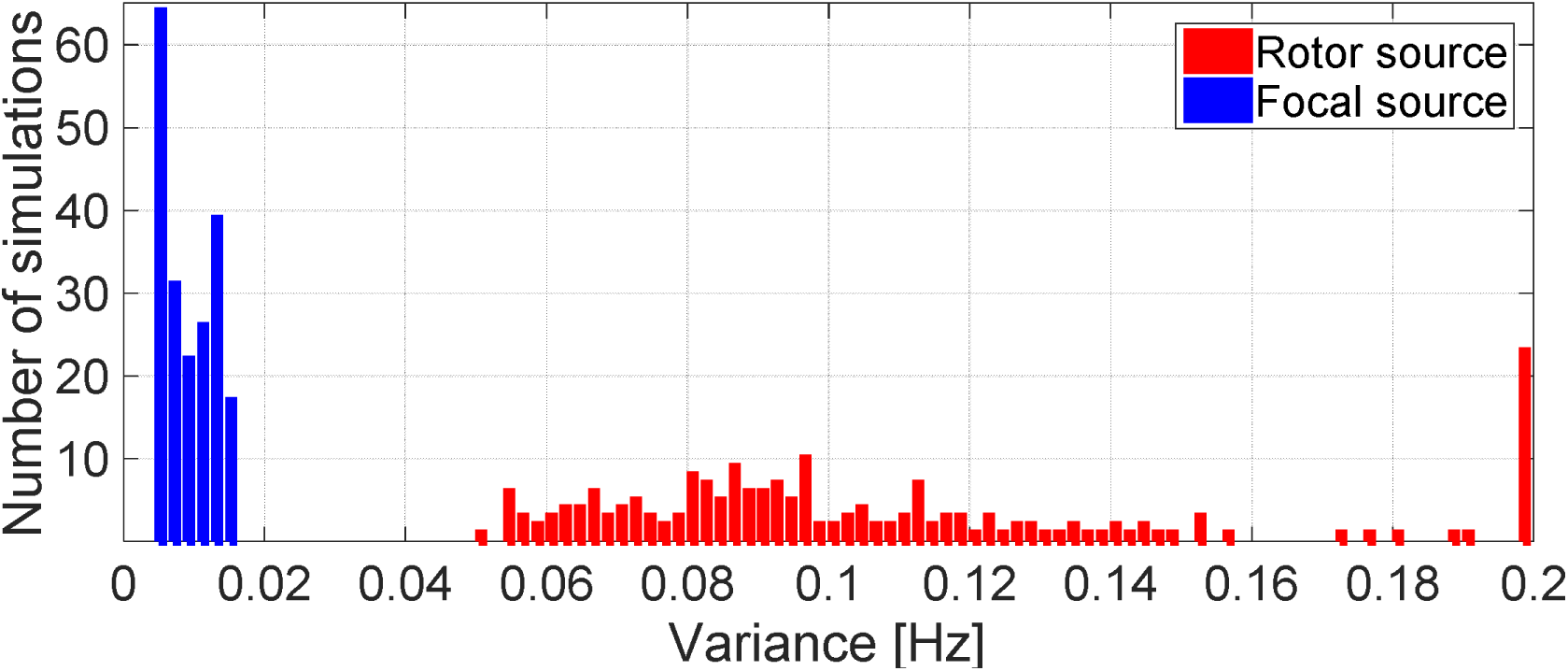
Histogram of the variance of the frequency vector for each simulation, for both source types: spiral wave (red) and ectopic foci (blue). This histogram clearly reveals two separate ranges for the expected variance for each source type and thus produces a well-defined threshold between 0.016-0.05 Hz for use in a binary decision.

**Fig. 10.**
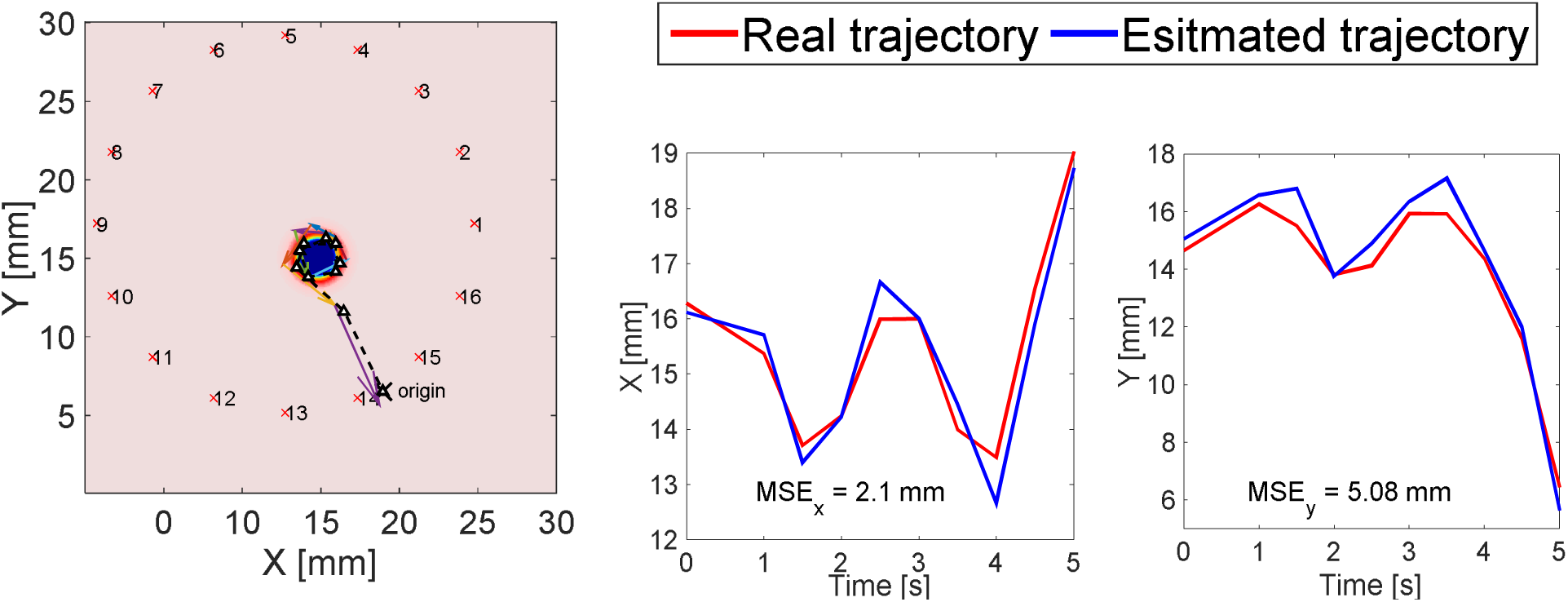
Reconstruction of the drifted spiral wave trajectory for a spiral wave with an origin at [19 mm, 16.5 mm] and an electrode array center at [13 mm, 13 mm]. All graphics are as shown in Fig. 3. The total MSE is 5.5 mm, and the distance between the final estimated point and the original rotor origin is 0.9 mm (left). Comparison of the original (red) and the reconstructed (blue) trajectories for both the X and Y directions reveals a similar pattern and results in small MSEs as indicated within the plots (right).

**Fig. 11.**
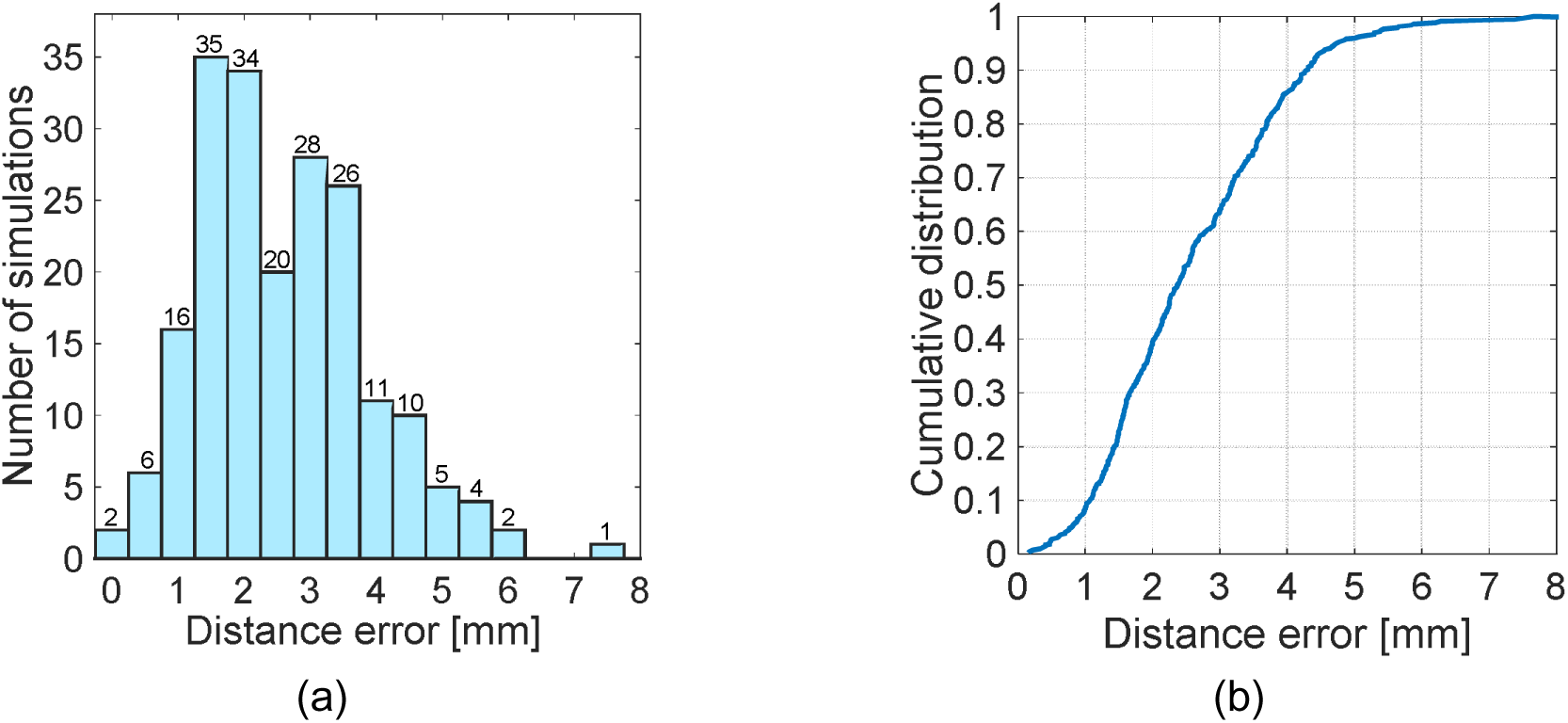
Analysis of the origin point prediction by the algorithm in random settings. Histogram of the distance error; i.e., the difference between the real origin point and the estimated point, for all 200 spiral wave simulations with random locations of origin points and electrode arrays. This diagram exhibits a Gaussian distribution with *μ* = 2.6 *mm*, *σ*^2^ = 1.6 *mm* (left) (a). In addition, a cumulative distribution function was generated to quantify the algorithm performance in arbitrary conditions. This revealed that almost 90% of the simulations resulted in a prediction of no greater than 4.3 mm from the original rotor origin point (b).

The histogram has a Gaussian-like distribution with μ=2.6mm and σ^^^2=1.6mm. In addition, all the origin points were detected at a distance that was not longer than 5.8 mm from the real origin point, excluding one simulation that yielded a distance of 7.3 mm.

Fig. 11(b) displays the cumulative distribution function of the error distance based on this histogram. As can be seen, most of the random rotor simulations had a small error distance; e.g., 50% of the simulations resulted in a 2.36 mm error and 90% of them had only 4.26 mm. Although the correct coordinates of the rotor origin were not determined with precision, and in fact the origin point is not a single coordinate in the tissue that can be pointed to, the algorithm was certainly oriented to the correct rotor origin area. Overall, these results demonstrate the capability of the method to detect tissue regions of electrical activity from a spiral wave, and to distinguish between reentry and focal activity.

## IV. Discussion

In this study, we investigated the feasibility of a Doppler-based algorithm to provide a new approach for robust detection of cardiac electrical conduction abnormalities. More precisely, we aimed at improved mapping and detection capabilities during atrial ablation procedures of two arrhythmogenic targets, spiral waves and ectopic foci, and developed a tool to differentiate between these two sources. Direct ablation of the spiral wave core or pivot point has been proven valuable for successful ablation and improved clinical outcomes, e.g., in acute AF patients [31]. However, common spiral wave mapping techniques during guided ablation are limited in this context since the rotor core cannot be determined with sufficient resolution. For instance, DF analysis is only able to detect the broader areas of highest DF in which the spiral wave exists, but these areas may include the majority of the atrial tissue, making the pivot point of the rotor and its periphery indistinguishable. Phase mapping is considered the gold standard for determination of the source type. While it is true that this is one of the most common methods for this determination, it has its own drawbacks: its ability to correctly identify the rotor pivot regions is limited by activation times that distort these maps, and it suffers from misleading phase and noise interference. Likewise, recent irregularity techniques (e.g. the irregularity index or CFAE) target the periphery of a source region, and may not necessarily identify the sites of the rotor tip or core [32]. Relatively new mapping methods based on signal processing (such as PCA [13], kurtosis and spatial Shannon entropy measurement [14]) are still being tested and have not been fully proven as more effective for the discovery of rotor pivot locations. Moreover, none of these mapping methods has been successfully implemented in clinical settings; hence, the need for another approach to improve mapping.

Here we proposed a new approach to rotor characterization and detection using spatial temperature gradients. To the best of our knowledge, the idea to exploit the notion that rotors drift towards low excitability regions in order to artificially attract and anchor rotors for potential ablation applications is novel. We consider that the interaction between rotors and temperature gradients can be utilized to help track rotors. Our numerical modeling results indicate that this utilization is workable by application of local temperature perturbations while measuring the electrical activity at several pre-set locations. The algorithm proposed here measures the Doppler shifts that arise from the rotor drift at various locations, and applies a back-track of the drift trajectory, which ends in the target origin area of the spiral wave. Hence, as supported by Gray [23], rotor drifting is a necessary condition to achieve Doppler shifts between signals. One of the mechanisms that causes the rotor to drift is related to heterogeneity in tissue excitability, which can be the result of gradients in the ionic channels [19]–[22]. However, inducing such gradients artificially is not feasible clinically. In contrast, applying a local temperature perturbation to the tissue via an inserted catheter, as we attempted to simulate, is simple, can lead to the desired gradient in excitability, and can initiate a controlled and known drift. In other words, by applying a local temperature perturbation, a gradient in the tissue excitability level is established, and this gradient leads to a controlled spiral wave drift, which creates Doppler shifts that can be manipulated for analyzing the arrhythmogenic source type.

Our algorithm results show that the reconstructed drift trajectory has the same pattern as the real one, including identical phases, similar peak-to-peak amplitudes and minimal MSE [see Fig. 3(d)]. In 90% of the 200 simulations conducted with a random location of the rotor pivot, the rotor origin point was estimated to be at a distance of less than 4.3 mm from the original one, as shown in Fig. 11. Given the fact that the rotor pivot is not a single coordinate but rather a small group of cells, these results may be interpreted as the detection of the rotor origin area within the tissue the physician should target during the ablation procedure.

The algorithm was tested as a function of several parameters. We showed that using more electrodes can reduce the total MSE exponentially and achieves better outcomes. Nonetheless, as shown in Eq. 13 and in Fig. 4(a), using 20 electrodes is sufficient for a minimal MSE, and increasing the number of measurements will not affect the algorithm results in terms of MSE. Application of the algorithm in noisy conditions yielded decreasing monotonic behavior as a function of the SNR and the number of electrodes [Fig. 4(b-c)]. Accordingly, employing the algorithm in a noisy environment with a sufficient number of electrodes should not affect the MSE to a large extent, certainly not enough to change the results significantly. While the noise effect is minimal, but still reasonable in these circumstances, future improvements of the system should include software and hardware filters and other noise reduction methods to diminish the noise influence completely.

We also found a relationship between the position of the electrode array and the location of the rotor origin and the temperature perturbation [Fig. 5–6]. When the center of the array was situated relatively close to the drift line, valuable results were achieved, both in terms of MSE and detection of the origin point. However, when the electrodes were shifted to the far periphery relative to the drift line, the Doppler effect was less noticeable in the electrodes and reconstruction was more difficult. Consequently, the ability to detect spiral wave was restricted and was shown to depend on the degree of proximity of the electrodes to the drift line. However, in the majority of cases the area in which the rotor was created was properly identified, which was the original goal of the method. One of our future ideas to overcome this hurdle is to use an automatically moveable temperature perturbation. In other words, by employing the perturbation at a certain point and analyzing the recorded signals in real time, the software will be able to decide whether to move it to another location, based on the obtained electrograms. Thus, after several iterations, we believe that the optimum position for the perturbation can be found so that the reconstruction will have a minimal error.

Our algorithm addresses the crucial need to distinguish between focal and reentry activity, two suspected sources of AF initiation. Measuring the activation frequency of a stationary rotor using several electrodes results in a constant value in all of them, similar to the frequencies obtained in the presence of a focal source. However, as we showed here, if the spiral wave drifts to a certain area, a Doppler effect occurs, and the activation frequencies are modified as a function of the location of the electrodes. We showed that focal activity, unlike a spiral wave, is not affected at all by the presence of a peripheral temperature perturbation, since the ectopic cells do not “sense” the temperature perturbation, such that the variance in the activation frequencies in the recorded signals can be neglected. Our results showed that the application of temperature perturbations during focal activity does not yield any source drift; hence the Doppler effect will not appear in the recorded signals. As a result, the variance of the frequency vector should not be more than 0.02Hz, as found in our simulations, and a binary decision can be made correctly, as was achieved in all 400 random simulations in this study.

Several limitations of our study should be noted. The ionic model (CRN) and the 2D tissue geometry model used here were considered to be simple and isotropic, without manifestations of complex AF conduction patterns such as spiral wave breakup or multiple wavelets. Nevertheless, the purpose of this study was to introduce a new approach for rotor detection. Our intent was to put forward a simplified, isotropic anatomy model in which a single rotor is active, in order to avoid any effects of geometric complexities that will make the analysis overly complex and increase the computational load. Future studies should include AF propagation patterns and realistic atrium anatomy to test the algorithm performance in more detailed scenarios and clinically realistic setups. Finally, this new approach requires a modification of the ablation catheter design that has a small probe for delivering the temperature perturbation. This should not be a major engineering problem since the typically used cryoballoon system already implements cold liquid as an energy source for AF ablation procedures [33]. While the integration of the temperature probe with the RF ablation systems is not standard practice in electrogram guided atrial ablation procedures, new catheters with these configurations can be designed, or, alternatively, merged with existing multielectrode lasso or flower catheters. This algorithm can also be used in the presence of other tissue heterogeneities that generate rotor drifting due to ionic mechanisms. Attempts to reconstruct its trajectory should be investigated to improve the mapping even without the use of the temperature gradient.

Nevertheless, this new method has advantages that outweigh its disadvantages. The full calculation takes only a few seconds, and can be made in real time during the ablation procedure. The results can be visualized, both in a quiescent tissue and in a tissue that sustains spiral wave activity or focal activity. It only need a small number of electrodes and is compatible with existing equipment. Finally, the estimated region of the rotor origin can be identified at high resolution, compared to other methods such as DF and CFAE, and can thus direct the clinical team to the target ablation site. However, it should be recalled that this is a proof of concept study, and further comprehensively investigated numerical simulations, in vivo experiments and clinical studies are needed before this technique can be translated into clinical practice. Our hope is that the mechanistic insights from this study can be utilized in the future for the design of a new methodology for AF characterization and termination, and that this study can contribute to the improvement of existing atrial ablation mapping capabilities, thus increasing success rates and optimizing arrhythmia management.

## Acknowledgment

The authors gratefully acknowledge Dr. Sharon Zlochiver for his helpful suggestions and comments for his article.

